# Assessing the potential of plains zebra to maintain African horse sickness in the Western Cape Province, South Africa

**DOI:** 10.1101/751453

**Authors:** Thibaud Porphyre, John D. Grewar

## Abstract

African horse sickness (AHS) is a disease of equids that results in a non-tariff barrier to the trade of live equids from affected countries. AHS is endemic in South Africa except for a controlled area in the Western Cape Province (WCP) where sporadic outbreaks have occurred in the past 2 decades. There is potential that the presence of zebra populations, thought to be the natural reservoir hosts for AHS, in the WCP could maintain AHS virus circulation in the area and act as a year-round source of infection for horses. However, it remains unclear whether the epidemiology or the ecological conditions present in the WCP would enable persistent circulation of AHS in the local zebra populations.

Here we developed a hybrid deterministic-stochastic vector-host compartmental model of AHS transmission in plains zebra (*Equus quagga*), where host populations are age- and sex-structured and for which population and AHS transmission dynamics are modulated by rainfall and temperature conditions. Using this model, we showed that populations of plains zebra present in the WCP are not sufficiently large for AHS introduction events to become endemic and that coastal populations of zebra need to be >2500 individuals for AHS to persist >2 years, even if zebras are infectious for more than 50 days. AHS cannot become endemic in the coastal population of the WCP unless the zebra population involves at least 50,000 individuals. Finally, inland populations of plains zebra in the WCP may represent a risk for AHS to persist but would require populations of at least 500 zebras or show unrealistic duration of infectiousness for AHS introduction events to become endemic.

Our results provide evidence that the risk of AHS persistence from a single introduction event in a given plains zebra population in the WCP is extremely low and it is unlikely to represent a long-term source of infection for local horses.

## Introduction

African horse sickness (AHS) is a viral disease of equids caused by the African horse sickness virus (AHSV), a RNA virus from the genus *Orbivirus*. The virus is principally transmitted by *Culicoides* spp. midges and is endemic in most sub-Saharan Africa with all nine types occurring regularly, affecting domestic horses, donkeys and wild equids, such as zebra. Infections in domestic horses result in significant clinical disease, leading to a high mortality rate, ranging from 50% to 95%, in naive animals, whereas infections in zebra are asymptomatic [1].

The presence of AHS in South Africa has resulted in a non-tariff barrier to directly trade live domestic equids with the European Union [2, 3]. In order to establish the trade of live equids from the country, an AHS controlled area was established within the Western Cape Province (WCP) in 1997. This controlled area, located at the very south-western tip of South Africa, consists of an inner AHS free zone, a surveillance zone and a protection zone which differ in their risk profile and the nature of the equine population in each zone [4]. The AHS controlled area was established in the WCP principally on the basis that the distance to the Kruger National Park (KNP), located in the north-eastern extremity of the country, would be large enough to effectively isolate local horses from the large zebra population present in KNP and in which AHSV circulates persistently [5], thereby lowering the risk of AHS introduction through seasonal spread. Furthermore, widespread vaccination of domestic horse populations outside the AHS controlled area is regularly implemented to further mitigate the risk of disease introduction in the AHS controlled area (Animal Diseases Act 1984).

Even though vaccinating domestic horses has markedly decreased the annual incidence in South Africa compared to pre-vaccination levels [6], AHS still remains endemic in most of the country, with approximately 500 cases reported annually in domestic equids [2]. In contrast, only sporadic outbreaks have occurred within the AHS controlled area, mostly linked to the re-assortment and/or reversion to virulence of the live attenuated AHS vaccine in use in the country [7].

AHS has not been reported in zebra since at least 1993 in the Western Cape Province (DAFF disease database) and albeit limited, prevalence survey results have not detected sero-positive animals in this area [6]. However, the risk of AHSV persisting in the WCP in the event that it is introduced in the local zebra population is a source of concern for the horse industry. Besides the increased cost in regaining free status if wild equids were infected, the geographic expansion of several midge-borne diseases and the emergence of arboviruses in Europe, causing large scale epidemics in livestock, have resulted in a sensitisation of European countries to the reality that the (re-)emergence of these viruses is a possibility, particularly for regions that have resident populations of *Culicoides* vectors [1, 8–10]. Such a sensitisation has highlighted the need to investigate zebra as a potential wildlife reservoir in the WCP to facilitate the re-establishment of trade in live equids from South Africa to Europe. Such a concern is further exacerbated by the fact that the most recent documented incursion of AHS into Europe (in 1987) was thought to have originated from southern Africa (Namibia) *via* the importation of zebra [1, 11]. An evaluation of the potential for zebra populations to act as a reservoir for the disease in the AHS control zones also ensures that established surveillance strategies, movement and other AHS controls remain relevant the current control activities in place.

In this study, we used a well described zebra population data-set within the WCP and including the AHS controlled area, along with published parameters relating to the populations and dynamics of AHS transmission, to inform an age- and sex-structured host-vector model and evaluate the likelihood that introduction events in the WCP would result in persistent AHSV circulation within a zebra population, similar to what is observed in KNP. As such, we focused our analyses to plains zebra (*Equus quagga*, formerly *E. burchellii*), the most abundant wild equid in South Africa and in which AHS is endemic in KNP [5]. This work not only provides robust risk-based evidence for improving AHS preparedness and surveillance, but also represents an opportunity to answer questions that were asked over 20 years ago regarding the required size of a zebra population to maintain AHS infection in South Africa [6].

## Materials and methods

### Modelling framework

#### Vector-host compartmental model

We developed an hybrid deterministic-stochastic vector-host compartmental model, where the spread of AHSV in the host (i.e. plains zebra) population is stochastically modelled whereas the spread of AHSV in vectors is assumed deterministic. Here, we considered that the transmission process of AHSV in plains zebra is only driven by *Culicoides imicola*. Fig 1 illustrates how both vector and host populations are structured and describes each population in regard to the spread of AHSV in the population.

**Fig 1.**
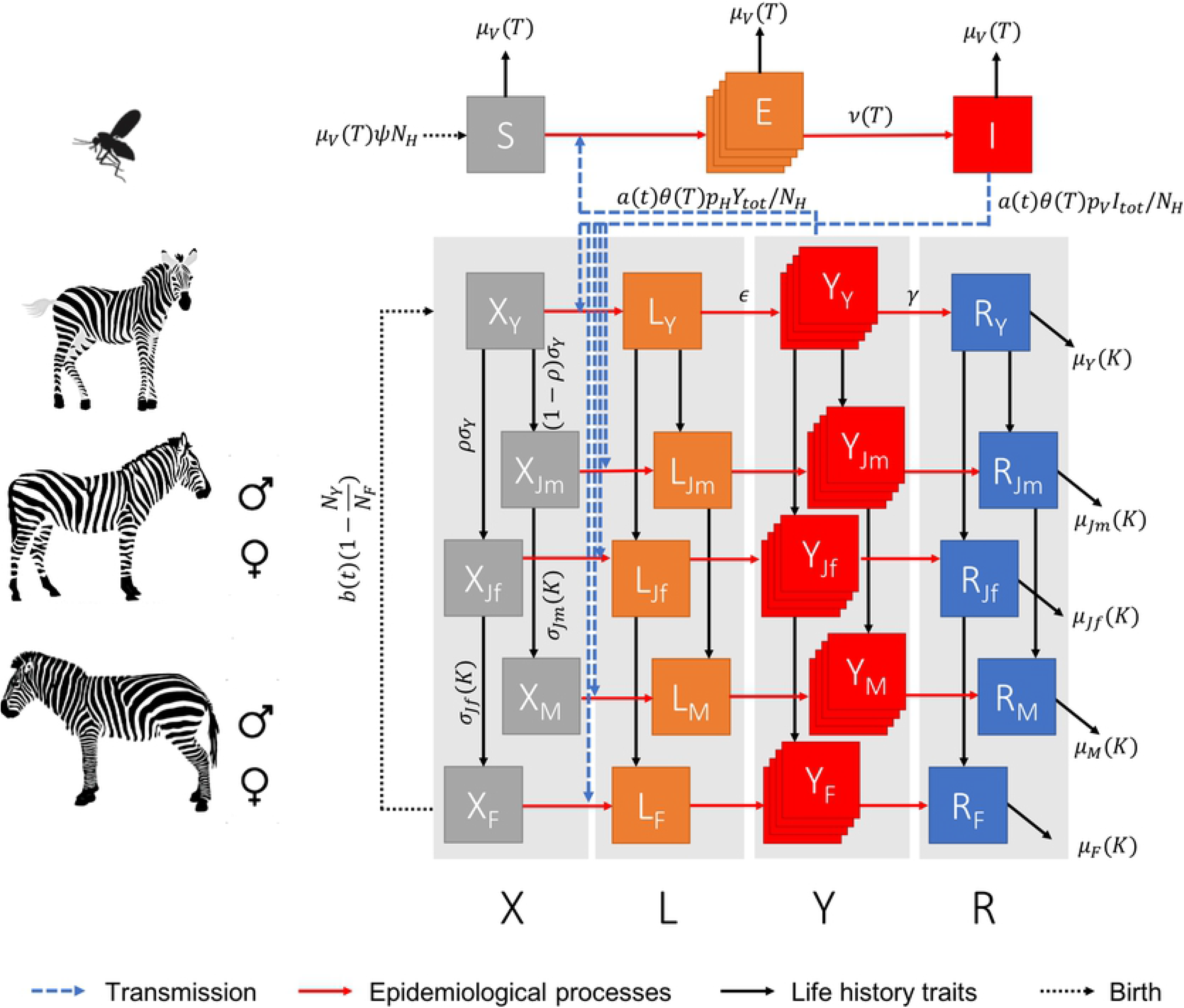
Schematic representation of the vector-host model. Zebra are either in a susceptible (*X*), latently infected (*L*), infectious (*Y*) or recovered (*R*) state, with subscripts *Y*, *Jf*, *Jm*, *F* or *M* indicating the age class of the animal, being foal, female juvenile, male juvenile, adult female and adult male, respectively. Vectors are either in a susceptible (*S*), latently infected (*E*) or infectious (*I*) state. The stacked squares denote that this compartment is divided into multiple stages. All rates between epidemic states and age classes are defined in Tables 1 & 2. However, some rates may depend on environmental conditions, denoted in brackets, with conditions being the daily mean temperature (*T*) recorded at sunset (i.e. between 18:00 and 21:00), the yearly carrying capacity (*K*) or the calendar month of the year (*t*). For clarity, rates were given only once per epidemic state or age class. Note also that the rate of mortality for each zebra age class would be constant for all epidemic states.

Our model shares common features with the AHS model of Backer & Nodelijk [12] and with the Schmallenberg and Bluetongue model of Bessell et al. [13] and Græsbøll et al. [14]. Particularly, a latent compartment for hosts has been included to account for the relatively long delay period between infection and infectiousness that is expected to occur in zebra as in horses. We also divided some compartments into five independent stages to better emulate transition processes and avoid exponential decays in both the incubation period of vectors and infectious period of hosts [12–14]. However, our model differs by (1) considering infection events in plains zebra will result in little to no mortality and will generate a life-long immune protection against a single AHS type, and (2) restricting our model to a closed population of zebra in a given holding, with a defined population size and surface area. The latter assumption was taken based on the large distance separating holdings with zebra in the WCP (mean = 204 *km*, SD = 118, 95% range 23 – 447 *km*) and the little recorded movements of animals between holdings. In addition to these assumptions, our model further differs by integrating the long-term population dynamics of the host population, separating the host population into five age classes. For simplicity, hosts are first considered born as a foal (*Y*, <1 year-old), irrespective of their gender, but then mature into either male (*J*_*m*_) or female (*J*_*f*_) juveniles when one year old. Male and female juveniles then mature into male (*M*) and female (*F*) adults (>3.4 year-old), respectively.

**Table 1.**
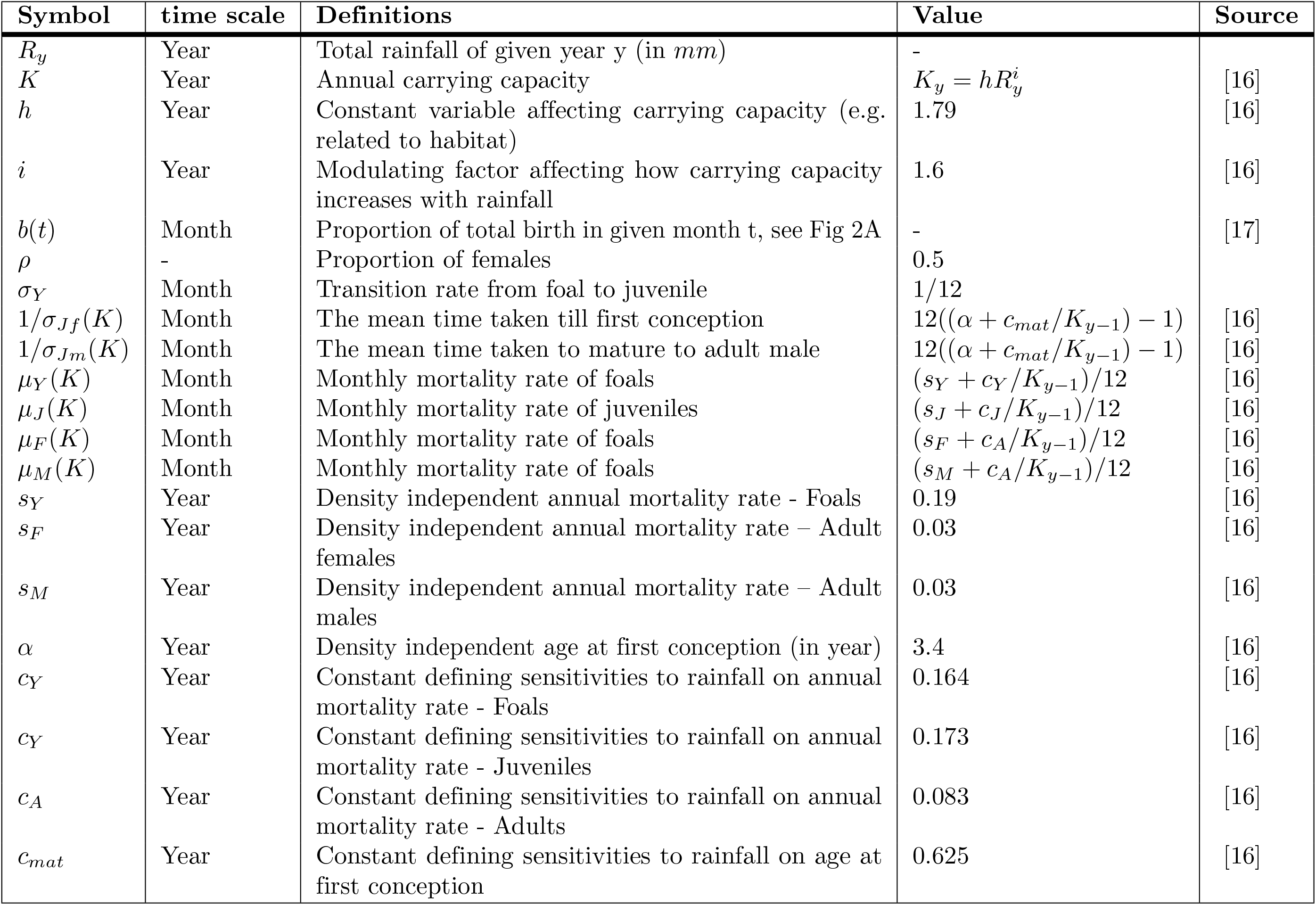
Parameters used in the zebra population dynamic model and their default values.

| Symbol | time scale | Definitions | Value | Source |
| --- | --- | --- | --- | --- |
| $R_y$ | Year | Total rainfall of given year $y$ (in $mm$) | - | |
| $K$ | Year | Annual carrying capacity | $K_y = hR_y^i$ | [16] |
| $h$ | Year | Constant variable affecting carrying capacity (e.g. related to habitat) | 1.79 | [16] |
| $i$ | Year | Modulating factor affecting how carrying capacity increases with rainfall | 1.6 | [16] |
| $b(t)$ | Month | Proportion of total birth in given month $t$, see Fig 2A | - | [17] |
| $\rho$ | - | Proportion of females | 0.5 | |
| $\sigma_Y$ | Month | Transition rate from foal to juvenile | 1/12 | |
| $1/\sigma_{Jf}(K)$ | Month | The mean time taken till first conception | $12((\alpha + c_{mat}/K_{y-1}) - 1)$ | [16] |
| $1/\sigma_{Jm}(K)$ | Month | The mean time taken to mature to adult male | $12((\alpha + c_{mat}/K_{y-1}) - 1)$ | [16] |
| $\mu_Y(K)$ | Month | Monthly mortality rate of foals | $(s_Y + c_Y/K_{y-1})/12$ | [16] |
| $\mu_J(K)$ | Month | Monthly mortality rate of juveniles | $(s_J + c_J/K_{y-1})/12$ | [16] |
| $\mu_F(K)$ | Month | Monthly mortality rate of foals | $(s_F + c_A/K_{y-1})/12$ | [16] |
| $\mu_M(K)$ | Month | Monthly mortality rate of foals | $(s_M + c_A/K_{y-1})/12$ | [16] |
| $s_Y$ | Year | Density independent annual mortality rate - Foals | 0.19 | [16] |
| $s_F$ | Year | Density independent annual mortality rate – Adult females | 0.03 | [16] |
| $s_M$ | Year | Density independent annual mortality rate – Adult males | 0.03 | [16] |
| $\alpha$ | Year | Density independent age at first conception (in year) | 3.4 | [16] |
| $c_Y$ | Year | Constant defining sensitivities to rainfall on annual mortality rate - Foals | 0.164 | [16] |
| $c_Y$ | Year | Constant defining sensitivities to rainfall on annual mortality rate - Juveniles | 0.173 | [16] |
| $c_A$ | Year | Constant defining sensitivities to rainfall on annual mortality rate - Adults | 0.083 | [16] |
| $c_{mat}$ | Year | Constant defining sensitivities to rainfall on age at first conception | 0.625 | [16] |

Briefly, we assumed that a host of the age and sex class *a* ∈ {*Y, J*_*f*_, *J*_*m*_, *F, M* can be either susceptible (*X*_*a*_), latently infected (*L*_*a*_), infectious (*Y*_*a*_) or recovered (*R*_*a*_). In the model, the exact number of hosts that change state or age class is determined by a binomial distribution with the probability *p*_*i*_ = 1 − *exp*(−*r*_*i*_Δ*t*), where *r*_*i*_ are rates of changes computed for each *i*^*th*^ transition and Δ*t* is the time-step. The model further assumes that no maternal immunity would be acquired by foals, leaving them susceptible to infection from birth. Similarly, no cross protection between AHS types was assumed; hence, all zebra are susceptible to any given AHS type, irrespective of historical exposure to other types.

In the model, we considered that adult female *C. imicola* were the only vector life stage involved in AHS transmission dynamics. We therefore discarded the fine population dynamics of vectors and the impact of environmental factors on eggs, larvae and nymphs, and assumed vectors would be subject to a constant hazard of dying. Vectors can either be susceptible (*S*), latently infected (*E*, i.e. in the extrinsic incubation period) or infectious (*I*); as such it considers that vectors will be infectious till their death. However, to keep the ratio between the number of vectors and the number of hosts *ψ* constant over time [13], we considered the vector population would replenished by the birth of susceptible vectors and fixed the total size of the vector population *N*_*V*_ such as *N*_*V*_ = *ψN*_*H*_ where *N*_*H*_ is the total size of the host population. The latter assumes, however, that the maximum number of vectors feeding on a single zebra in each day is constant between hosts, irrespective of their age class.

It is worth noting that, while the epidemiological processes of AHS in both zebra and vector populations are computed at a daily time-step, the population dynamics for zebra is modelled at a monthly time-step. As such, in each given month, the epidemiological process would be carried out daily over a constant population.

The virus can be transmitted from an infectious vector to a susceptible host at a rate depending on the number of infectious vectors *I*, the biting rate *θ*(*T*) and the activity rate *a*(*t*) which adjusts for the expected number of midges on host at time *t*. As such, the product *a*(*t*)*θ*(*T*)*I* gives the number of infectious bites per day [13]. However, not all bites will be infectious to any given host; hence, we defined the probability *p*_*V*_ that the bite of an infectious vector on a susceptible host is successful in transmitting the virus. In the model, the number of susceptible hosts of any given age class that would be infected by infected midges is taken from a binomial distribution *Bin*(*X*_*a*_, Λ), such as Λ = 1 − exp(−*λ*_*V*_) and where *λ*_*V*_ is the transmission rate from vector to host and is defined as:

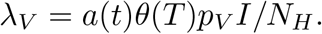

This assumes that the probability that an infectious vector bites a susceptible host is equal to the fraction of susceptible individuals of a given age class in the host population *X*_*a*_/*N*_*H*_ [12].

Similarly, the transmission rate from an infectious host to a susceptible vector is derived from the number of infectious bites per day *a*(*t*)*θ*(*T*)*I*, the probability *p*_*H*_ that the bite of a susceptible vector on an infectious host is successful in transmitting the virus [12, 13]. However, we assumed that all infectious hosts have the same ability to infect vectors, irrespective of their age class. The probability that vectors bite an infectious host is therefore the fraction *Y*_*tot*_*/N*_*H*_, where 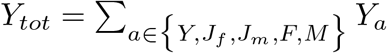 is the total number of infectious hosts in the population. Consequently, the transmission rate from host to vectors is defined as:

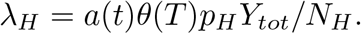

All other parameters used in the model and their default values are defined in Tables 1 and 2. Particularly, the biting rate *θ*(*T*), the extrinsic incubation rate 1*/ν*(*T*) and the mortality rate *μ*_*V*_ (*T*) were considered depending on the temperature *T* (in °C) as midges will bite more often and the virus will replicate faster in vectors, requiring less time to be transmissible, when temperatures increases. However, with increasing temperatures, vectors will also die sooner. These relations were all extracted from [12].

**Table 2.**
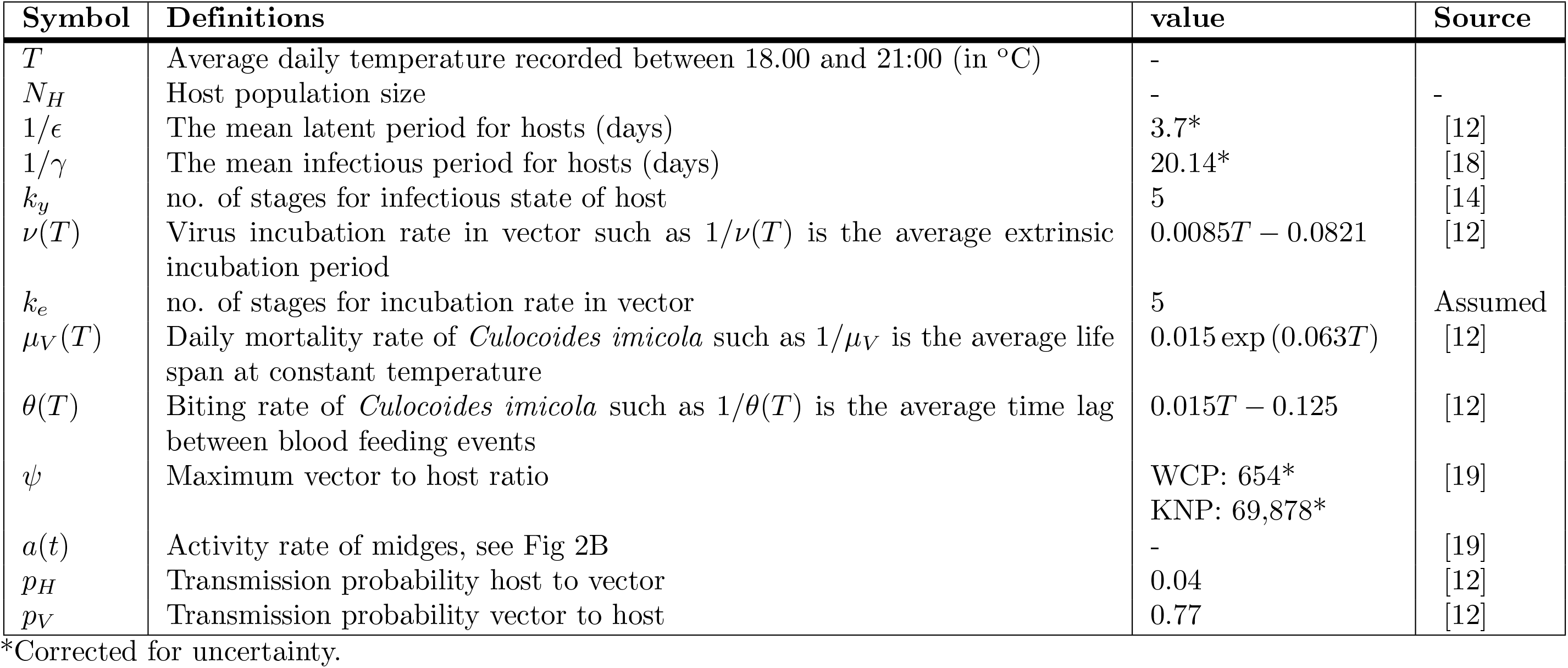
Parameters used in the AHS transmission model and their default values.

| Symbol | Definitions | value | Source |
| --- | --- | --- | --- |
| $T$ | Average daily temperature recorded between 18.00 and 21:00 (in °C) | - | - |
| $N_H$ | Host population size | - | - |
| $1/\epsilon$ | The mean latent period for hosts (days) | 3.7* | [12] |
| $1/\gamma$ | The mean infectious period for hosts (days) | 20.14* | [18] |
| $k_y$ | no. of stages for infectious state of host | 5 | [14] |
| $\nu(T)$ | Virus incubation rate in vector such as $1/\nu(T)$ is the average extrinsic incubation period | $0.0085T - 0.0821$ | [12] |
| $k_e$ | no. of stages for incubation rate in vector | 5 | Assumed |
| $\mu_V(T)$ | Daily mortality rate of *Culicoides imicola* such as $1/\mu_V$ is the average life span at constant temperature | $0.015 \exp(0.063T)$ | [12] |
| $\theta(T)$ | Biting rate of *Culicoides imicola* such as $1/\theta(T)$ is the average time lag between blood feeding events | $0.015T - 0.125$ | [12] |
| $\psi$ | Maximum vector to host ratio | WCP: 654*<br>KNP: 69,878* | [19] |
| $a(t)$ | Activity rate of midges, see Fig 2B | - | [19] |
| $p_H$ | Transmission probability host to vector | 0.04 | [12] |
| $p_V$ | Transmission probability vector to host | 0.77 | [12] |

\*Corrected for uncertainty.

The model was developed in the programming software R version 3.4.4 [15].

#### Zebra population dynamics

We based our model on the age- and sex-structured population dynamic model of a plains zebra population in Laikipia District, Kenya, developed by Georgiadis et al. [16].

Briefly, the model adjusts vital rates and age at first reproduction through a rainfall-mediated density-dependent (RMDD) mechanism. This RMDD modulates vital rates by a factor that depends on the ratio of carrying capacity *K*_*y*_ to population size *N*_*H*_. Here, the carrying capacity is defined as the maximum number of individuals that could be supported in a given year, at maximum reproductive output and was estimated as a function of the total rainfall (*R*_*y*_, in *mm*) recorded in the location of interest over an entire year *y* such as 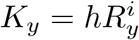, where *h* and *i* are constant variables.

While the original model was designed and fitted over annual records of abundance of plains zebra, we adapted this modelling framework to account the temporal variations of births within each given year. As such, numbers within each age and sex class were calculated in successive monthly time-steps as shown in Fig 1. While foal numbers for each given year were calculated as equal to the number of adult females surviving from the previous year and that did not give birth in the previous year, the number of foals born in each given month varies as a function of the monthly density of foals *b*(*t*) (Fig 2A). The birth function *b*(*t*) was derived from empirical observations in the proportion of <1-month-old foals for two populations of plains zebra in the KNP between August 1969 and March 1973 [17]. Foals that mature to juveniles are then allocated to each sex, assuming foal sex ratio to be unity and identical survival. Female and male juveniles will then mature to adulthood at a rate *σ*_*Jf*_ (*K*_*y*_) and *σ*_*Jm*_(*K*_*y*_), respectively. Foals, juveniles, adult females and adult males will all die at a different monthly rate, i.e. *μ*_*Y*_ (*K*), *μ*_*J*_ (*K*), *μ*_*F*_ (*K*) and *μ*_*M*_ (*K*), which also depend on the carrying capacity *K*_*y*_.

**Fig 2.**
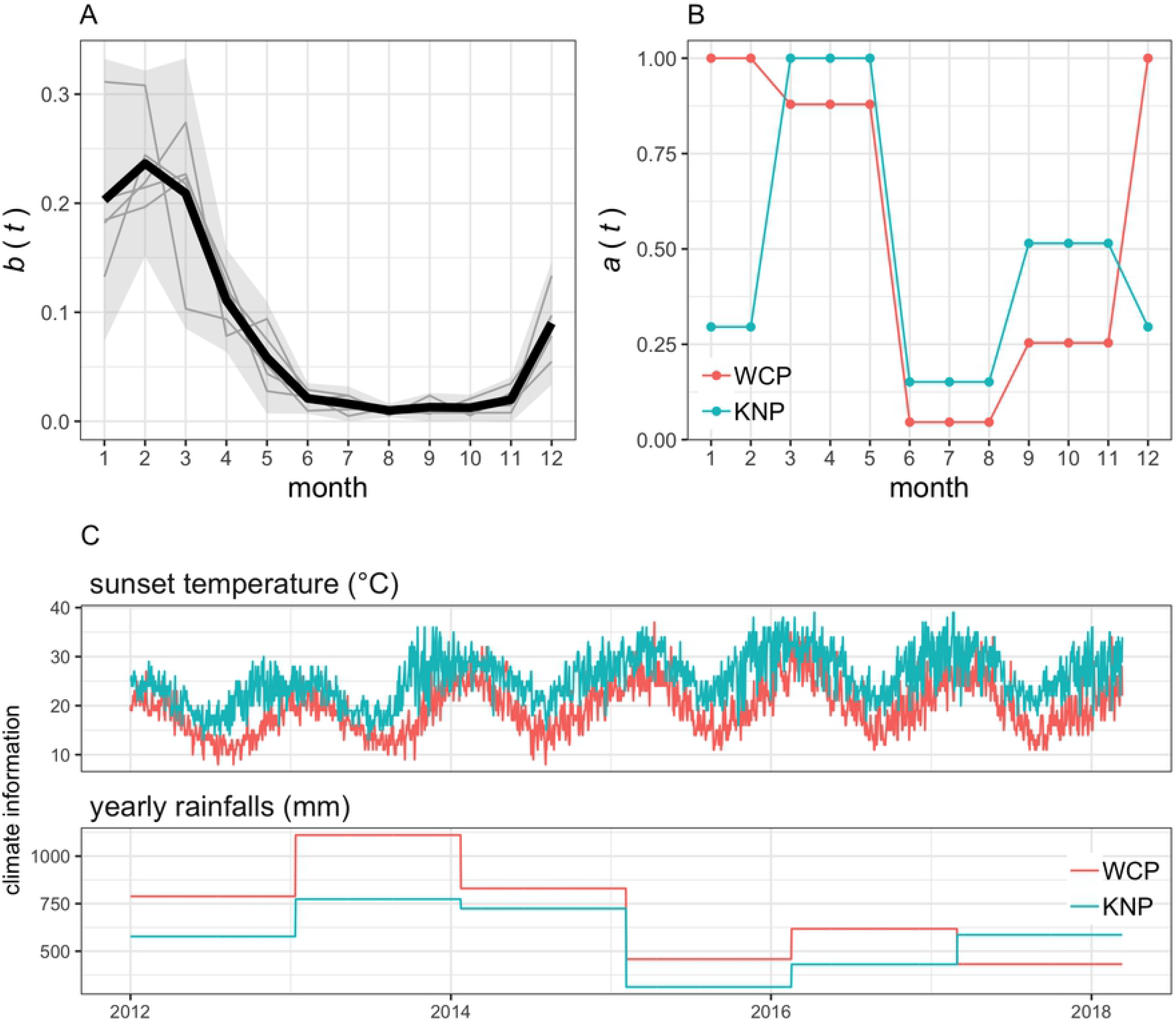
Time-related functions and variables used in the model. (A) The zebra birth function *b*(*t*) as defined by the monthly density of the annual number of births of plains zebra in the Kruger National Park (KNP) between August 1969 and March 1973. Solid grey lines show the adjusted birth data used to calculate *b*(*t*) and extracted from Smuts [17] who reported the temporal variation in the proportion of <1-month-old foals for two populations of KNP. Black solid line and grey envelop shows the birth function *b*(*t*) and its 95% confidence interval. Note that 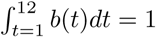. (B) The activity function *a*(*t*) of *Culicoides imicola* as defined by the proportion of the maximum number of *C. imicola* per hosts per day recorded in each month in the Western Cape Province (WCP) and KNP. Data extracted from Venter et al [19]. Trap catch records from Stellenbosch and Eiland (see Fig 3A for location) were used to characterise midge activity in the WCP and KNP, respectively. (C) Daily variations in sunset temperature and variations of the annual rainfalls recorded in Stellenbosch (WCP) and Eiland (KNP).

All vital rates and parameters used to model the zebra population dynamics were fitted from empirical observations and related to changes in annual records of rainfall in Laikipia District [16]. While we acknowledge that the ecological conditions in the Laikipia District and our study sites are markedly different, we considered that using site-specific rainfall data would control for such differences. As such, parameters controlling the zebra population dynamics were kept similar to those fitted in [16], though modified to account for monthly time-steps.

#### Estimating epidemiological parameters and accounting for uncertainties

Parameters driving the population dynamics of zebra and those drawn experimentally were well defined in Georgiadis et al. [16] and Backer & Nodelijk [12], respectively. We therefore considered that natural variations for these processes would be accounted for by the stochasticity of the system. On the other hand, particular attention was required to estimate and account for the uncertainty on some epidemiological parameters given the low level of understanding of the epidemiology of AHS in zebra. In particular, three parameters were perceived uncertain or unknown and could significantly affect the outcomes of our analyses. These were the mean latent period in zebra (1*/ϵ*), the mean infectious period in zebra (1*/γ*) and the maximum vector to host ratio (*ψ*). While different strategies were used to account for the uncertainty in each parameter, they all consisted in randomly drawing parameter values from defined distributions at the start of each simulation.

The estimate the latent period of infected hosts (1*/ϵ*) was drawn from a Gamma distribution with a mean of 3.7 days and a standard deviation of 0.9 days (corresponding to 16 latent stages). This is similar to what has been done by Backer & Nodelijk [12] and considering that the latent period in zebra would be similar to what was recorded experimentally from five unvaccinated challenged horses [20–22]. This is however consistent with infection outcomes from seven experimentally challenged plains zebra [18] showing that virus isolation from blood yielded positive results on day four.

Given that it was assumed that zebra would be infectious for a period following a Gamma distribution with five infectious stages (consistent with models of bluetongue virus spread [14]), we only had to account for the uncertainty in the mean infectious period (1*/γ*). To estimate the mean infectious period of plains zebra, infection outcomes from seven experimentally challenged plains zebra [18] were used. These seven zebra were last viraemic at 20, 13, 17, 24, 24, 24, 40 dpi, which, after accounting for a conservative 3-day latent period, yields an infectious period of 10 to 37 days. Using these data, the mean infectious period in zebra was estimated at 20.14 days, which would lead to zebra being infectious for a median of 18.8 days and a 95% range interval of 6.5 – 41.3 days. We further account for the uncertainty in the mean infectious period by randomly sampling its estimate at the start of each simulation from a normal distribution with a mean of 20.14 days and a standard deviation of 2 days, i.e. a 2.5% – 97.5% interval of 16.2 – 24.1 days.

The maximum number of midges per plains zebra (*ψ*) were extracted from records of seasonal abundance in *Culicoides* populations (including *C. imicola*) in the vicinity of livestock at seven sites of South Africa between October 1983 and December 1986 [19]. For the purpose of this study, records of only two of the seven sites were used (Fig 3A): Stellenbosch (33°56’S, 18°52’E) and Eiland (23°40’S, 30°45’E); and were chosen to characterise the seasonal midges’ activity in the WCP and in KNP, respectively. Venter et al. [19] reported the total number of collection events and the total number of *Culicoides* trapped per season (spring, summer, autumn and winter) as well as the proportion of *C. imicola* over the total number of *Culicoides* collected. Estimates of *ψ*, together with the activity rate *a*(*t*), were then computed from these data assuming that one light trap collection using downdraft blacklight traps mimics, when placed near host, the daily host seeking activity of *Culicoides* toward one individual zebra as well as its seasonal variation [12]. We therefore translated trap data directly to vector to host ratios, identified the maximum number of *C. imicola* per collection events to estimate *ψ* and adjusted remaining records to compute *a*(*t*), with monthly values of *a*(*t*) ranging from 0 to 1 (Fig 2B). We further account for the uncertainty in the maximum number of midges per plains zebra by randomly sampling the estimate of *ψ* from a Poisson distribution at the start of each simulation.

**Fig 3.**
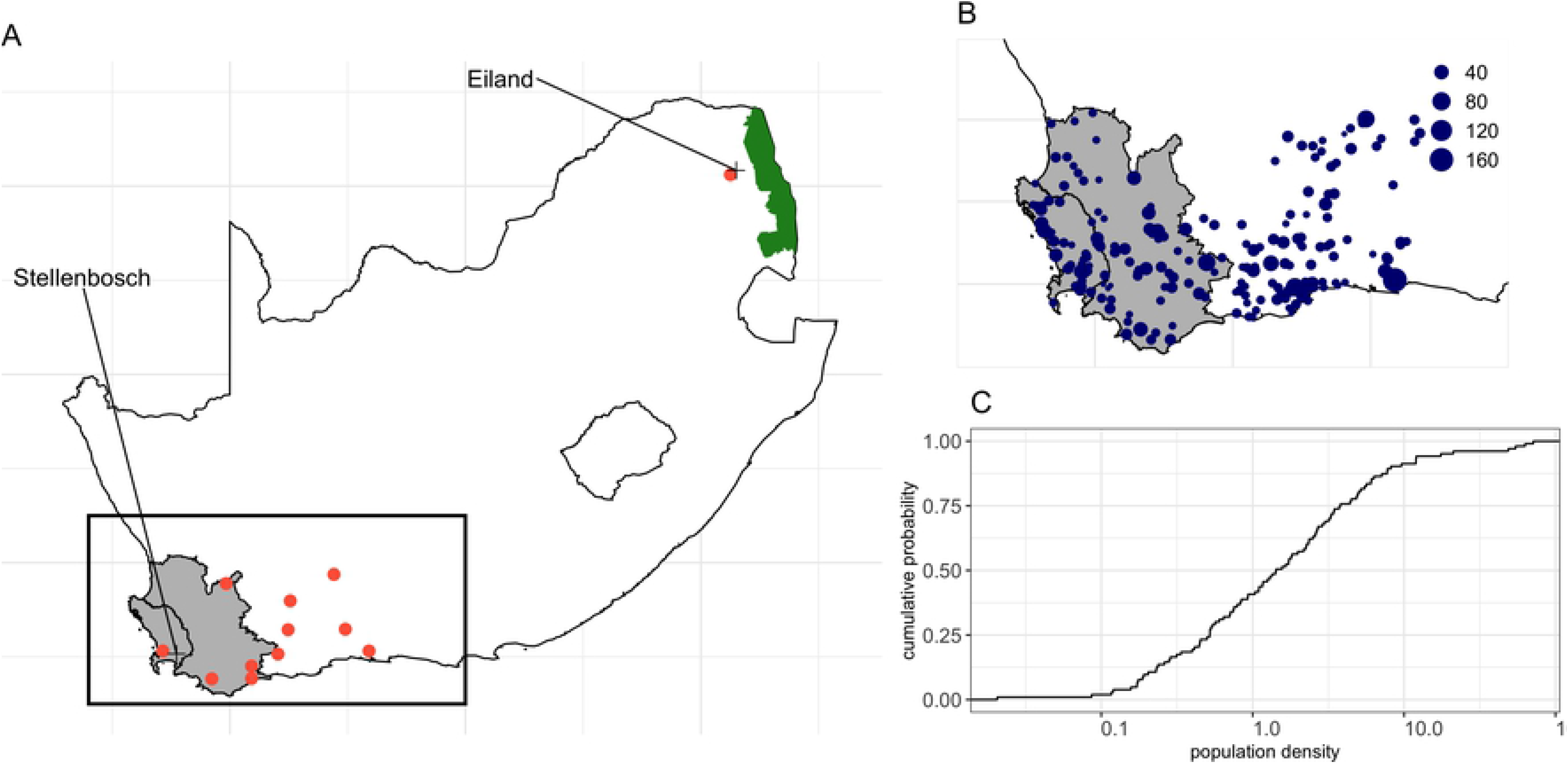
Study area and holdings with a reported population of plains zebra. (A) Location of the 12 sites in which climate data were extracted from. Red solid dots indicate sites at which climate information were downloaded, whereas crosses indicate the location for which midge abundance data were reported in Venter et al. [19]. Rectangle represents the extent of the study area. (B) Location of the 212 holdings with at least 1 reported plains zebra that were considered in this study. Size of each dots are indicative of the size of the zebra population. (C) Distribution of the population density of the 102 holdings with at least 1 reported plains zebra and a known surface area. Grey shaded areas in A and B represent the AHS controlled area of the Western Cape Province, while the green shaded area in A is the Kruger National Park (KNP). Note that the *x*-axis in C is log_10_-scaled.

### Model implementation

Through this study, 40 independent incursions of AHS were tested for each individual scenario considered. Incursion events were assumed resulting from the introduction of a single infected (but not infectious) adult female into a given plains zebra population. Unless otherwise stated, the introduction event would occur on the first day of the study period (1st January 2012) and disease would be left to spread for six years (or 2192 days).

Because records of population structure in each considered population are missing, we assumed that each population of plains zebra have a similar population structure at the start of the simulations as found in Georgiadis et al. [16]. As such, at the start of each simulation, the population included 16% foals, 25% juveniles, 24% adult males and 35% adult females, with a general male-to-female ratio of 0.7.

In this study, we considered AHS reached an endemic state if the virus is still actively present in the population after 2000 days (i.e. 5.5 years). We also computed various statistics over the tested 40 independent incursions: namely, the median (and its interquartile range, IQR) and the maximum duration of the outbreak as well as the probability of outbreak lasting >1 year, >2 years and >3 years.

### Study sites

The compared study areas are in the Western Cape Province of South Africa within which is the AHS controlled area (greyed area as shown in the rectangle in Fig 1A), and the KNP, which is located in the Limpopo and Mpumalanga Provinces in the north-eastern part of the country (Fig 1A). To our knowledge AHS has not been reported in zebra since at least 1993 in the Western Cape Province (DAFF disease database) and albeit limited, prevalence survey results have not detected sero-positive animals in this area either [6]. In contrast, prevalence surveys in various locations in the north-eastern parts of the country have shown a high sero-prevalence for AHS in zebra, with close to 100% of juvenile or adult zebra testing positive to at least one AHS serotype [5, 6]. While it is considered that endemic circulation in the KNP takes place there is no comprehensive data, to our knowledge, available regarding the incidence rate of this circulation.

The difference in AHS occurrence in the different parts of South Africa is not only defined by an existing wild populations that may maintain ongoing circulation, but the environmental conditions conducive to transmission which impact host, vector and virus [6]. While the KNP is vast and consists of multiple climatic zones, it is characterised by a hot wet summer and a lack of winter frost which has been associated with the *Culicoides* vector being present to facilitate year-round transmission [23]. The Western Cape in contrast, and particularly the AHS controlled area, has a Mediterranean climate with cold wet winters with associated frost and hot dry summers.

### Scenario tested

To explore how initial conditions of the AHS introduction in plains zebra may affect the risk of AHS to remain circulating for an extended period of time in the WCP, we implemented two scenarios of disease introduction.

First, AHSV was introduced in the largest population of plains zebra recorded in the WCP and on different dates. Here, we considered that introduction events occurred on the first day of each month for four years (i.e. from January 2012 to December 2015). Simulation were then left running until January 2018.

Secondly, we focused on the situation where AHSV was introduced during the high risk period that was identified in the previous scenario. In this scenario, AHSV will be then introduced in a given holding where plains zebra was reported. In this scenario, we considered that introduction events occurred on the first day of the earliest month showing the highest risk of AHS persistence. All holdings were tested individually.

It is important to note that, although the introduction of the infection may occur at different times and on different holdings, all simulations were kept starting on the first day of the study period, providing consistent dynamics of the zebra population.

### Sensitivity analysis

We assessed the sensitivity of the risk of AHS persistence with respect to variations in (1) the population size of plains zebra in which AHS is introduced into, and (2) the mean infectious period, 1*/γ*. Not only would these help to evaluate the sensitivity of the model outcomes to the uncertainty related to disease incursions and the epidemiology of AHS in zebra, it will also would clarify the thresholds at which AHS would constantly persist and become endemic in the WCP.

In each situation, we considered that AHS would be introduced during the high-risk period previously identified and in a holding located in the control zone and for which climatic conditions are typical to the situation in the WCP. Here, we considered that the zebra population in which AHS was introduced would remain at a constant population density equal to the average population density reported in the WCP (i.e. 1.5 plains zebra per *km*^2^).

To investigate the impact of the population size on the risk of AHS persistence, we considered AHS to be introduced in holdings of various population sizes, varying from 5 to 50,000 individuals. To investigate the impact of the mean duration of infectiousness of plains zebra on the risk of AHS persistence, we progressively increased the value of 1*/γ* from 20 days to 50 days. However, because the impact of increasing 1*/γ* may inherently depend on the size of the zebra population in which the virus is introduced, we further evaluated the sensitivity of the model to variations of 1*/γ* for four population size (10, 50, 100 and 150 individuals).

### Data

#### Climate data

Climate data were downloaded from www.worldweatheronline.com using their historical weather application programming interface (API), through R using package *rvest* [24]. The API allows searching for data using longitude and latitude for given spatial locations. Data were requested for 11 randomly spatially sampled locations in the greater WCP and 1 location representative to the climate and midge ecology in the KNP. Fig 3A shows the location of the 12 sites in which climate information were downloaded. Daily data from 01 January 2012 to 01 January 2018 were extracted, and included the maximum, minimum temperature (in °C) as well as the three-hourly average temperature (in °C) and the three-hourly total precipitation (in *mm*).

Under field conditions in South Africa, and as measured with light traps, *C. imicola* are most active immediately after sunset, whereas host-seeking activity remain low the rest of the day [25]. Similarly, annual rainfalls are the main driver of population for the plains zebra population in Laikipia District, Kenya [16]. As such, yearly records of rainfall and the daily mean temperature recorded at sunset (i.e., between 18:00 and 21:00) were considered the only drivers of vector and host populations and were computed from the extracted climate data.

For the purpose of this work, vital traits of plains zebras and *C. imicola* population in each considered location in the WCP were driven by the climate records (i.e. yearly rainfalls and sunset temperature) of the nearest location for which data were downloaded, respectively. In contrast, simulations for KNP were always driven by the same set of temperature and rainfall data. Fig 2C compares the daily variations of sunset temperature and the yearly precipitation for Eiland and Stellenbosch as exemplars of the climate variables in KNP and the WCP, respectively.

#### Population data

In South Africa, it is mandatory to register properties that keep zebra in the AHS controlled area (South African Animal Diseases Act, Act 35 of 1984). Under a data sharing agreement, records of the location, surface area, population size and observation date for all holdings that keep plains zebra in the WCP were provided by CapeNature (www.capenature.co.za). Overall, 2498 plains zebras were reported in the WCP, all spread across 212 holdings. Fig 3B shows the location and size of the populations of plains zebra kept in the WCP. Most of the populations of plains zebra in the WCP are small, with a median population size of 7 zebra (IQR: 4 – 14) and a maximum of 168 plains zebra.

Over the 212 holdings with at least 1 reported plains zebra, surface areas of 110 could not be retrieved. We therefore imputed the unknown surface areas (in *km*^2^) at the start of each simulated incursion events by first (1) randomly drawing the population density (in individual*.km*^−2^) from the 102 holdings with a known surface area (Fig 3C), and (2) converting the population density into a surface area as a function of the reported number of zebra. Plains zebra in the WCP are kept at a median density of 1.50 individual per *km*^2^ (IQR: 0.52 – 3.78) and a maximum of 71.4 individual per *km*^2^.

KNP is one of the largest game reserves in Africa and covers an area of 19,485 *km*^2^. Kruger National Park has around 70% of the free-roaming population in South Africa with 29,000 – 43,450 individuals in 2012 [26]. To the best of our knowledge this is the only South African population that is migratory and, while zebra are likely to show preference for different vegetation zones within the park (www.krugerpark.co.za/Kruger_National_Park_Regions-travel/kruger-park-eco-system.html) we assumed that the land in KNP was fully available to them, suggesting that the population density of zebra in KNP ranged between 1.49 and 2.23 zebra per *km*^2^. For simplicity, the plains zebra population in KNP was considered at similar population density as in the WCP, with a population size of 30,000 individuals and a density of 1.54 zebra per *km*^2^.

## Results

### Baseline simulations

Fig 4 provides a comparison on how the abundance of the plains zebra populations in KNP and in the WCP may vary during the study period. It is clear that these two populations behave differently. To facilitate comparison, we provide a projection of the population size considering a 10% annual population growth (the solid red line in Figs 4A & 4C). The population in KNP showed little stochastic variations in abundance, with an initial increase in population with an annual rate of population growth of 4.2% (95% CI 4.1% – 12.5%), reaching a maximum of 36,672 individuals. From January 2015, however, the population of KNP declined progressively due to poor availability of forage from successive years of low precipitations. In contrast, the population in the WCP showed high variability in predicted abundance, with high chances for significant reduction due to unsustainable population densities. Such a marked variability is partly due to the fact that the surface area of this population was unknown and therefore imputed from known population densities in the WCP. However, in simulations with population density of <3.77 individuals per *km*^2^, the population will increase with a mean annual rate of 4.8% (IQR: 2.9% – 6.6%), reaching a max of 235 animals.

**Fig 4.**
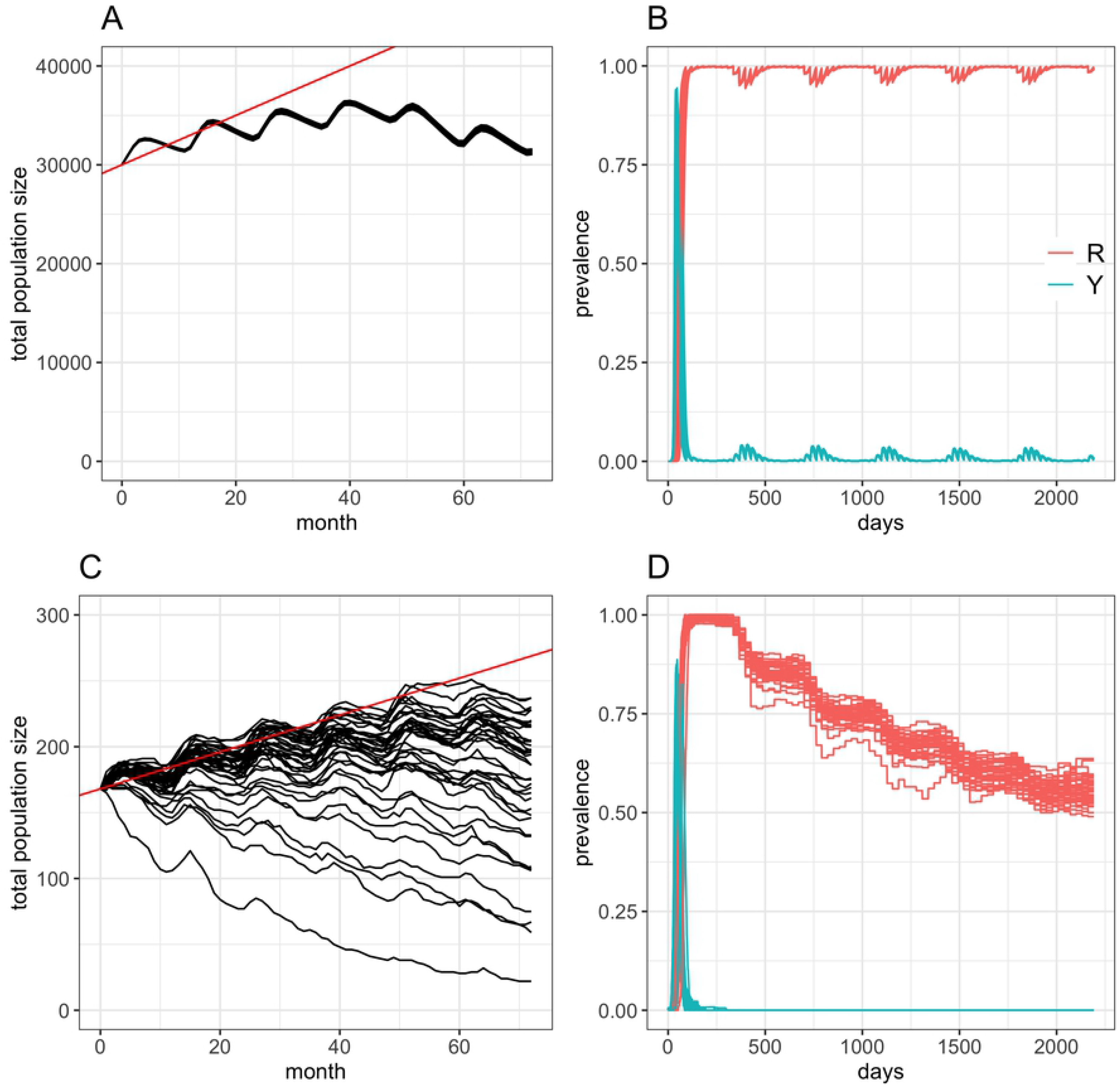
Baseline dynamics of the plains zebra population and AHS prevalence after introduction at the start of the study period in KNP and the WCP. (A-B) Temporal evolution of the predicted number of plains zebras (A) and those that are in the infectious and immune states (B) following 40 independent incursion events occurring on January 1st 2012 in the KNP plains zebra population. (C-D) Temporal evolution of the predicted number of plains zebras (C) and those that are in the infectious and immune states (D) following 40 independent incursion events occurring on January 1st 2012 in the WCP plains zebra population. We assumed that the KNP population of plains zebra (A and B) included 30,000 individuals at the start of each simulation. Similarly, we considered incursions in the WCP (C and D) occurred in the largest population of plains zebra reported in the area (N=168) at the start of each simulation. Solid red lines in A and C are the 10% annual increase in population size. Note the difference of scales in the y-axis.

In KNP, introducing an infected AHS adult female in the zebra population at the start of the study period resulted in AHS becoming endemic in 100% of the 40 simulations tested (Fig 4B) but at a low level of prevalence. One year post introduction, i.e. when an endemic equilibrium is reached, 0.99% (95% CI 0.62% – 1.36%) of zebras would be infected either at the exposed (*L*) or infectious (*Y*) stages, whereas 98.9% (95% CI 98.6% – 99.4%) of zebra would be immune to AHS (i.e. in *R* state). S1 Fig shows the daily variations in the prevalence of infectious hosts stratified by age class. It is clear that the persistence of AHS in the plains zebra population in KNP is driven by foals, with a clear seasonality driven by their birth. In January and February, the mean monthly proportion of infectious foals is 4.2% to 4.5%, whereas <0.3% of the foals will be infectious from July to September. Over all simulations implemented in KNP, nearly all foals (>99%) will be immune to infection when reaching juvenile age.

In contrast, the introduction of an infected AHS adult female in the largest population of plains zebra in the WCP never resulted in an endemic situation (Fig 4D). Instead, AHS persisted in the population for a median period of 155 days (95% CI 147 –161) and never circulated for more than 1 year.

### Temporal variations in the risk of AHS persistence in the WCP

Table 3 and S2 Fig show how long AHS persists in the population for each month of the study period and for each of the 40 independent incursion events that were tested. Two main outcomes can be established from this scenario: (1) introducing AHS in April-May would have the greatest impact on virus persistence and would represent a high-risk period in the WCP, with AHS likely to remain circulating for more than a year in 37.7% (30.6% – 45.5%) and 34.4% (27.5% – 42.0%) of the time (Table 3); (2) even introduced in the high-risk period, AHS will not persist beyond 500 days post introduction.

Instead, introducing AHS in April will generate an outbreak which would last for a median of 193 days (IQR: 110.5 – 421 days). For comparison, outbreaks of AHS started between June and September would only last in plains zebra population for less than a month (Table 3, S2 Fig).

**Table 3.**
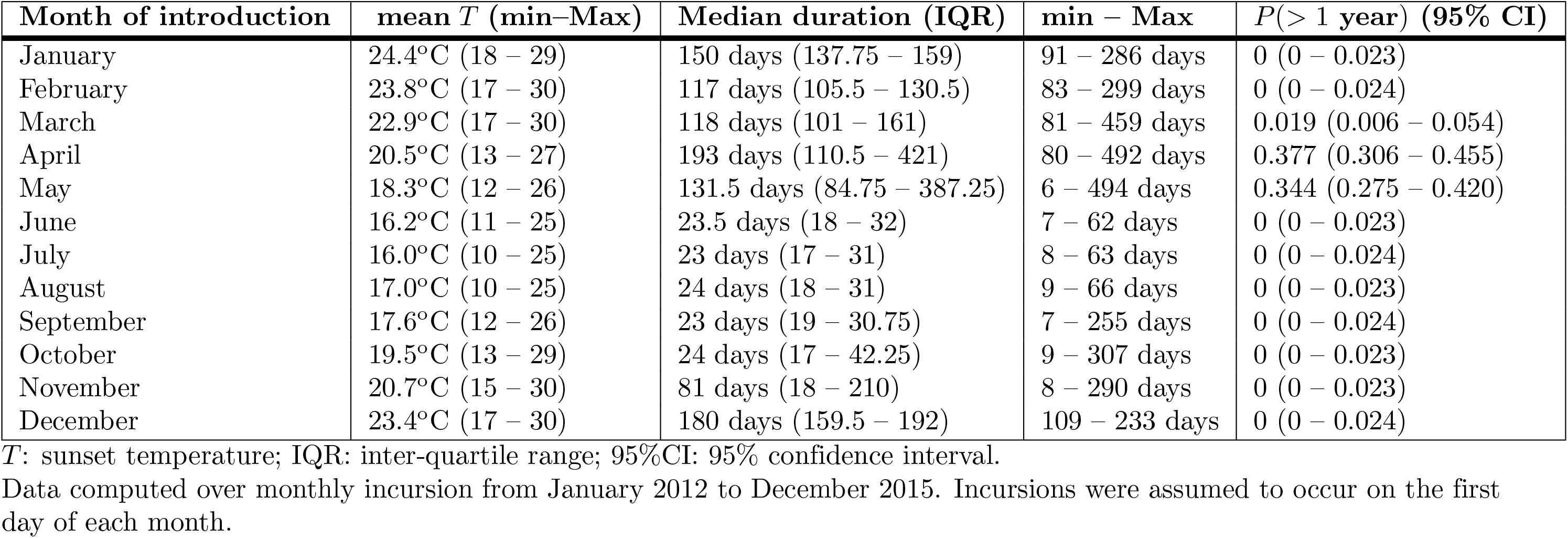
Risk of AHS persistence for the largest population of plains zebra in the WCP and stratified by the month of AHS introduction events.

### Spatial variations in the risk of AHS persistence in the WCP

We explored how conditions in each holding with plains zebra in the WCP may affect the risk of AHS persistence should introduction occur on April 1st, 2012.

Over all scenarios tested with regards to the incursion site, AHS outbreaks would last for a median of 135 days (IQR 123 - 161), with AHS circulating for less than 300 days in 94% of simulations (Fig 5A). Although AHS was never found persisting for more than 4 years in the WCP, 78, 5 and 1 holdings were able to generate outbreaks lasting >1 year, >2 years and >3 years, respectively. Among the 78 holdings that were able to keep AHS circulating for >1 year (denoted as high risk in Fig 5B), 13% and 26% generated outbreaks lasting for >1years in only 1 or 2 simulations over the 40 considered. Fig 5C & S3 Fig show the spatial distribution of the risk of long-lasting circulation of AHS in the WCP.

**Fig 5.**
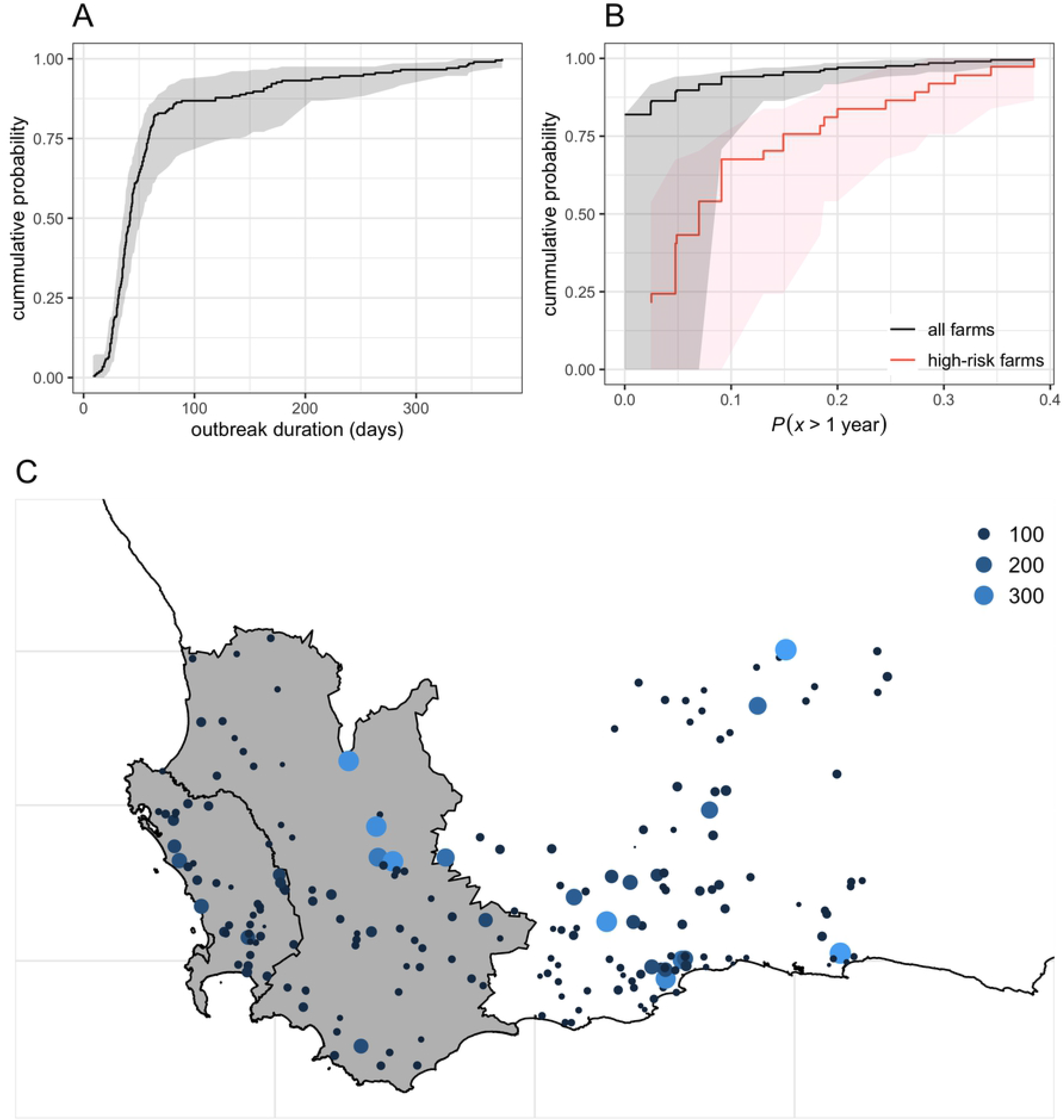
Distribution of the risk of AHS persistence among all individual holdings with plains zebra in the WCP. Cumulative probability distributions of the (A) median and IQR outbreak durations and (B) the probability that AHS outbreaks would persist for >1year computed over 40 independent AHS introductions in all individual holdings. (C) Spatial distribution of the risk of AHS persistence as measured by the median outbreak duration (in days) among holdings with plains zebra in the WCP. Here AHS was consistently introduced in April 2012.

To explore what conditions would allow long-lasting circulation of AHS in the WCP, we plotted the distribution of the median outbreak duration as a function of the size of the zebra population at the start of the simulation (Fig 6A). Two distinct groups of holdings may be observed with holdings denoted “A” showing orders of magnitude shorter median outbreak duration than those denoted “B”. These two groups are spatially defined (Fig 6B), with group A involving holdings along the coast of the WCP, whereas group B includes holdings located inland, on the plateau. Given the limited number of factors affecting the model, it is clear that differences in the dynamics of the temperature used to drive AHS epidemiology for each holding may affect the risk of AHS persisting in the WCP. S4 Fig shows the dynamics of the sunset temperature for all sites we used to extract climatic data that were driving holdings in group A and group B. Clearly, sunset temperatures in group B are warmer in summer (more often between 25°C and 30°C) and colder temperatures in winter (more often between 5°C and 10°C). While our model indicates that these temperatures enable *C. imicola* to survive and allow AHS to persist in the environment for 2 to 3 seasons after the initial outbreak period, this needs to be validated with entomological and epidemiological data. For the remaining of this study, we will therefore imply that these two groups of holdings are representative to two climatic zones and will denote them “climatic zone A” and “climatic zone B”.

**Fig 6.**
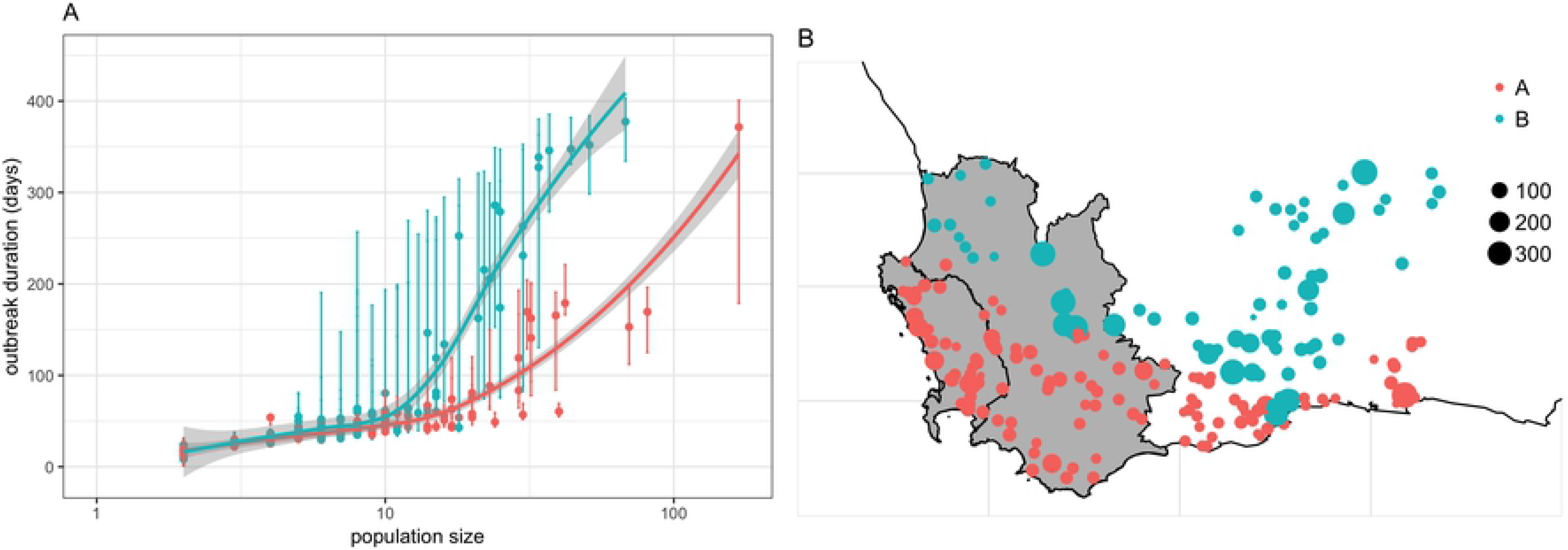
Distribution of the risk of AHS persistence as a function of the size of the plains zebra population in all individual holdings in the WCP. (A) Median, and IQR, duration of AHS outbreaks in all holdings considered in the WCP. This distribution shows two groups (groups “A” and “B”) of holdings differing from outcomes of AHS incursion events. (B) Spatial distribution of the risk of AHS persistence stratified by their group. Note that the large error bars in (A) highlight the large uncertainty for holdings that has no record of surface areas. Curves in (A) are the smoothed conditional means of the median outbreak duration. Size of dots in (B) represent the median outbreak duration (in days).

### Sensitivity analysis

Fig 7A and S5 Fig show the effect of increasing the size of the zebra population in which AHS is introduced on the outbreak duration and the likelihood of AHS persistence in the WCP. Clearly, conditions in climatic zone B are more likely to allow long term circulation of AHS in plains zebra than those in climatic zone A. In particular, incursion of AHS in zone A would seldom last until the end of our simulation period (i.e. for >5.5 years or >2000 days), whereas incursion in zone B would become endemic in a wide range of scenarios. For climatic zone A, only 3 simulations lasted for >2000 days, and required a population of plains zebra of at least 50,000 individuals. In contrast, AHS became endemic in at least one simulation for a population of at least 500 individuals in climatic zone B. Of interest, 1 in 4 incursions in climatic zone B would become endemic in a zebra population of 1500 individuals, whereas AHS would always persistently circulate if the population included 5000 individuals. All simulations would last at least 1 year for populations of at least 500 individuals and at least 250 individuals in holdings located in climatic zones A and B, respectively. Finally, a population of at least 500 and 1500 individuals would be required for holdings in climatic zone B to suffer outbreaks lasting >2 years and >3 years respectively for every incursion of AHS (S5 Fig). In contrast, at least a population of 25,000 and 50,000 zebra would be required for outbreaks lasting >2 years and >3 years respectively in 60% of the incursion events in holdings of climatic zone A.

**Fig 7.**
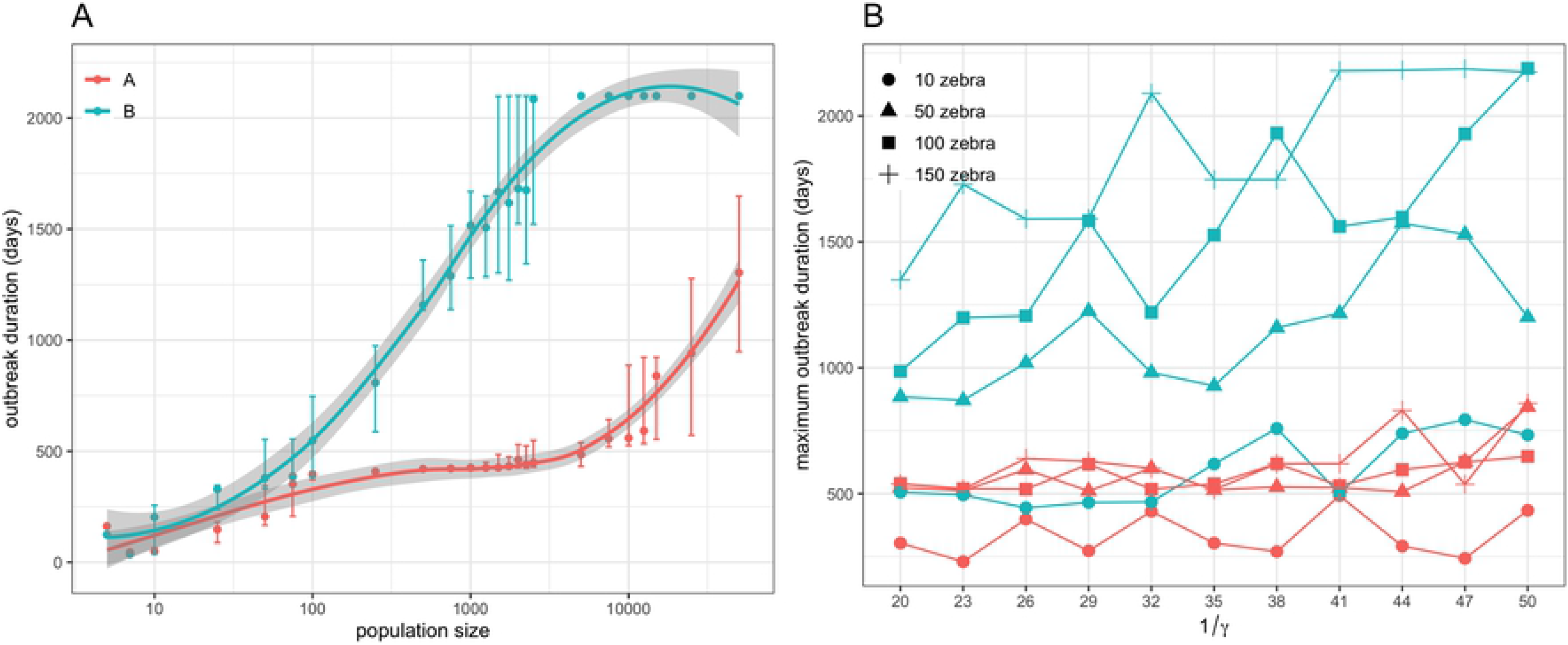
Sensitivity of the risk of AHS persistence to variations in population size and infectious period. (A) Effect of variations in the size of the plains zebra population on the risk of AHS persistence as measured by the median outbreak duration. (B) Effect of variations in the mean infectious period 1*/γ* for AHS in plains zebra on the risk of AHS persistence as measured by the maximum outbreak duration. In (B), changes in the risk of AHS persistence as a function of 1*/γ* was computed for various population sizes. Risk of AHS persistence was computed over 40 independent AHS introduction events that consistently occurred in April 2012. All considered zebra populations were set with a population density similar to the median density reported in the WCP. Here is shown the outcomes for incursions in a typical holding located in either climatic zone A or climatic zone B.

The effect of increasing the infectious period 1*/γ* for zebra on the risk of AHS persistence in the WCP is shown in Fig 7B and S6 Fig. Again, conditions in climatic zone A and B are markedly distinct. When incursion occurs in climatic zone A, outbreaks would never persist and would seldom last for >1 year, irrespective to the duration of 1*/γ*. In contrast, incursion occurring in climatic zone B may circulate for >3 years in a large enough zebra population (i.e. *≥*50 individuals). In particular, outbreaks can become endemic in climatic zone B for population of at least 100 individuals. However, for this to happen, zebra would need to be infectious for an extended mean duration of infectiousness. For example, an introduction event of AHS in a plains zebra population of 150 individuals lead to an endemic circulation of AHS in at least one simulation in climatic zone B when 1*/γ* ≥ 41 days, which, at population-level, would lead zebras being infectious for a median of 38.3 days and a 95% range interval of 13.3 – 84.0 days. It is however worth noting that, with such population level infectious period, the odds to observe the infectious period similar to those obtained from the experimental infection [18] is 1 over 30 trillion (*P* = 3.41 × 10^*−*14^). If the introduction were to occur in a population of 100 zebra, outbreaks would only become endemic if 1*/γ* =50 days, which, at population-level, would lead zebras being infectious for a median of 46.7 days and a 95% range interval of 16.2 – 102.4 days.

## Discussion

The results of our analyses on the transmission and circulation of AHS in zebras in South Africa provided an improved understanding of how AHSV spreads in this wildlife population. To do so, we combined and adapted two previously published modelling frameworks to account for the impact of the foaling season of plains zebra on the circulation of AHS between vectors and hosts. Using this model, we clarified conditions in which incursions would persist and become endemic in wild population of equids. Particularly, we showed that populations of plains zebra present in the WCP are not sufficiently large for AHS introduction events to become endemic and that coastal populations need to be >2500 individuals for AHS to persist >2 years, even if zebras are infectious for more than 50 days. Should AHSV be introduced in the coastal population of the WCP, AHS cannot become endemic unless the population involves at least 50,000 individuals. In contrast, inland populations of zebra in the WCP may represent a risk for AHS to persist if populations grow beyond 500 individuals.

Beyond the benefit of improved knowledge regarding the epidemiology of AHS in zebra in the WCP, our findings provide a basis to review and justify the current regulations from both South African Veterinary Services and international trade stakeholders to prevent introduction of AHSV in the wild and domestic equine populations of the WCP and, should an introduction occur, ensure the disease-free status is regained quickly. In our study, we found that the highest risk of prolonged circulation of AHS in zebra populations is likely to occur if AHSV is introduced in April, when sunset temperatures range from 13°C to 27°C (Table 3) and midges biting activity is high (Fig 2B). In contrast, lowest risk period was between June and September, where introduction events would rarely generate outbreaks lasting for more than a month. These findings consolidate animal movement regulations currently enforced for the AHS controlled area which only allow the import of zebra between June and August. In addition, they further support the recent amendment in the vaccination protocols [27] that strictly restricts the use of vaccination to the period between June and October to avoid AHSV outbreaks due to re-assortment and/or reversions to virulence of live attenuated vaccines [7]. Both of these measures are likely to mitigate the risk of introducing AHSV in zebra populations and ensure the virus would not remain circulating for extended periods within the controlled area.

This work was facilitated by accessing the register of holdings keeping zebra in which the location and the abundance of all zebra subspecies (plains zebra and others) were recorded. In South Africa, it is mandatory to register properties that keep zebra in the AHS controlled area (South African Animal Diseases Act, Act 35 of 1984). As such, we are confident that all populations of zebra currently present in the WCP are included in our study. However, we only have the records of holdings having zebra in 2018 and the total number of animals last counted. We therefore considered that abundance of zebra at the start of our simulation (January 2012) were at similar size as in recorded in the register and that population structure would be similar to [16]. We acknowledge that this creates some uncertainties regarding the number of zebra populations present as well as their abundance throughout the study period. While we believe that these assumptions make our results more conservative, they also highlight the need to keep regularly updated information on zebra populations. Regularly updated censuses would help to not only verify the veracity of our assumptions but also will allow ongoing validation and justification of the control measures in place for zebra in relation to AHS.

The WCP is divided into two climatic zones which produced vastly different epidemiology of AHS and risk of AHSV persistence in zebra. In our model, only sunset temperature was assumed modulating the transmission of AHS between zebra and vectors, thereby ignoring the impact of other factors on population dynamics and activity pattern. Particularly, we considered neither the suitability of the environment for breeding (soil type, water and species specific dung which may be important as a breeding substrate), nor the impact of freezing temperatures and daily humidity [28, 29]. While we believe these may not have significant impact for incursions in climatic zone A, discarding their effects have likely inflated vector winter survival estimates and ability for incursions to spread [23], and therefore resulted in overestimating the risk of AHS persistence for incursions in climatic zone B. Indeed, climatic zone B is mostly located in areas of plateau and inland where their arid climate is unsuitable for *C. imicola* [28]. Despite this, low records of abundance of *C. imicola*, as well as low prevalence level of AHSV in midges and equines, have been reported in Namaqualand [30] and in southern Namibia [31], both areas sharing similar climatic characteristics as climatic zone B. While we acknowledge that local irrigation practices may help maintaining suitable microclimate for *Culicoides* vectors in arid areas [28, 32], whether such a low circulation of AHSV is solely due to the few *C. imicola* present or together with other competent subspecies remain to be clarified. However, it seems unlikely that these may be sufficient to allow maintaining AHSV in plains zebra.

Given the predicted higher risk for plains zebra to maintain AHS in climatic zone B, there is a marked need for the quantitative characterization of the impact of ecological factors other than temperature (such as soil quality, humidity and rainfalls) on the survival, biting and breeding activities of *Culicoides* in South Africa. Unfortunately, entomological studies allowing reliable measurements suitable for use in epidemiological models are still scarce. In this context, the work in [25] should be commended and represents an exemplar toward better understanding environmental drivers in midges abundance and activities. A previous study in South Africa has shown that the minimum records in monthly values of both land surface temperature (LST) and normalized difference vegetation index (NDVI), an indicator of abundance of live green vegetation (hence, humidity), were the best predictors of the mean daily blacklight trap capture of *C. imicola* [33]. As such, coupling our model with predicted abundances from remote sensing information, similar to those in [33, 34], could represent a feasible alternative to improve the accuracy of our assessment in the future.

In this study, we focused on plains zebra as it is the zebra subspecies with the largest population in both the WCP and South Africa, and in which AHSV is endemic in KNP [5,6]. However, other subspecies are present in the WCP, notably the Cape Mountain zebra (*Equus zebra zebra*) and the Hartmann’s Mountain zebra (*Equus zebra hartmannae*). While the latter is present in very low numbers in the WCP (67 individuals in 4 herds), the population of Cape Mountain zebra is now (after many years of conservation efforts) at a similar size as that of plains zebra [35]. Furthermore, over the 1714 individuals present in the entire WCP (as recorded in the CapeNature database), nearly half of the Cape Mountain zebra population (*n* = 842 individuals) is concentrated in a single nature reserve, the Karoo National Park. This conservation land is in the north eastern region of the WCP, approximately 250 *km* from the border of the AHS controlled area and in the heart of the climatic zone B. Knowledge on the ecology and the epidemiology of AHS in Cape Mountain zebra is limited, but if they are similar to those in plains zebra, and in the unlikely event that climatic zone B unaccounted climatic characteristics are not detrimental, such a large population would maintain ongoing circulation of AHSV and increase the risk that AHS would be endemic in the WCP. Although our model could be modified to account for their ecology, this would require longitudinal data recording abundance and structure of their populations over long period of time. Until these are available, it will be difficult to evaluate if these zebra subspecies may represent greater risk of AHS persistence than plains zebra in the AHS controlled area of South Africa.

Despite being widely distributed, there is still a dearth of research and reliable data on the population dynamics of plains zebra in southern Africa. The population dynamics of plains zebra has however been previously characterised from >20 years of abundance records in Laikipia District, Kenya [16] and upon which a simulation model has been developed and validated [16, 36]. Not only does it represent a key tool to better manage plains zebra populations in Laikipia, such a model highlighted the role of annual rainfalls as a key limiting factor in the population growth of plains zebra. In this study, we capitalised on such a relationship and assumed that using annual rainfall information recorded in either the WCP or KNP would be sufficient to characterise the population dynamics of plains zebra in our study areas. Although our model behaves according to our beliefs and the anecdotal records of population growth in the area, it cannot be validated. Similarly, little is known on the level of AHS prevalence in plains zebra in South Africa apart from a single longitudinal prevalence study carried out in 1991/1992 [5] showing that almost 100% of the free-living zebra foals born in the KNP were sero-positive for at least one serotype of AHSV once reaching adulthood. In addition, a continuous transmission cycle of virus has been shown between *Culicoides* midges and zebras in the KNP [6]. While these are again consistent with our results, they only emphasise the critical need of systematic longitudinal population abundance and surveillance records in plains zebra to not only validate our assumptions and model structure, but also better clarify the role of zebra in sustaining AHSV in South Africa.

When developing our model, several assumptions were made to simplify the epidemiology of AHS in plains zebra and overcome the lack of information and recent data. Firstly, for simplicity, we assumed that all age classes are affected by AHS similarly. By doing so, however, we did not account for the potential role of maternal immunity of foals, previously reported [5] and included in early models of AHS in zebra [37]. Although our results are consistent to field observations [5, 6], the maintained circulation of AHS in KNP is driven by the seasonal dynamics in foal abundance (see S1 Fig). As such, not including the lapse of time during which foals are protected from infection could impact the timing of the peaks of AHS incidence. While it is unknown if this would markedly affect the potential for AHS incursions to become endemic in plains zebra, such an impact remains to be verified in the future.

We further considered that populations of plains zebra were isolated not only from other populations of wild or domestic equids, but also from other non-susceptible hosts such as cattle and small ruminants. We also assumed that *C. imicola* would behave similarly across all age- and sex-classes when choosing which host to feed upon. Both of these assumptions may affect the attractiveness and the availability of hosts to vectors. The availability of other host species, and the impact it may have on biting behaviour of vectors, has been shown critical in the spread of midge-borne diseases such as bluetongue and AHS [38–41]. Particularly, increases in abundance of host with limited susceptibility to AHS would dilute the potential for disease transmission as they represent additional opportunities for *Culocoides* midges to feed. This would depend on the attractiveness of a given host as well as its abundance and availability [41]. It is not yet clear how zebra would attract *C. imicola* if presented to multiple host species to feed on. However, given the long infectious period for AHVS in zebra, even a low attractiveness may represent a risk for disease transmission into domestic equid populations in WCP. Although it remains unclear if the presence of multiple host species would affect the potential of plains zebra to maintain AHS in WCP, they may also act as potential sentinels for AHSV circulation in zebra populations. In the context of post-outbreak surveillance, this can represent an alternative to actively sampling zebras which logistically difficult and associated with high cost.

Finally, we assumed that *C. imicola* will act as the only vector of AHS in zebra. Given that *C. imicola* is recognized as the most important vector of AHS in South African [23, 42], and that the understanding in the feeding behaviour of other *Culicoides* and their relation with zebra remain limited, this assumption was believed reasonable. In South Africa, however, at least 13 other *Culicoides* species, including *C. magnus*, *C. nevilli*, *C. zuluensis*, *C. bolitinos*, and *C. enderleini*, are known to be orally susceptible to infection with AHSV [43, 44]. Particularly, *C. bolitinos* is a proven vector of AHSV [45] and represents the most likely driver of AHS transmission in cooler mountainuous areas of South Africa [42, 46]. In WCP and KNP, *C. bolitinos* is however 10 to 100 times less abundant than *C. imicola* [47, 48]. Nevertheless, increasing the number of vectors species, particularly if their ecology and niche significantly differ, could increase the complexity of the transmission dynamics of AHSV [23] and increase the likelihood that disease would spread and reach endemic state [49]. Again, whether this endemic state could occur in zebra populations of the WCP despite relatively low population abundance should be considered in future research.

## Conclusion

While it has long been known that zebra, if present in large enough populations, play a role in the epidemiology of AHS in South Africa, it has not, until now, been explicitly established to what degree they could play a role in maintaining infection should they be associated with a sporadic outbreak in the WCP. While there is still much to be established regarding the host, vector, environmental and viral components of the disease in wild equids, we show that, in the current population structure (both herd and region-level), it is unlikely that zebra are in populations large enough to maintain a persistent AHS infection in and around the South African AHS controlled area. We have shown there is a potential for within-region differences in risk of persistence of AHS and further study is required to confirm this potential.

The current sensitisation of global stakeholders involved in animal trade and public health to the threat of Orbiviruses such as African horse sickness means that zebra, and particularly factors that might result in large intermingling populations, should not be ignored in control and surveillance planning both prior to, during and in the post-outbreak period in the Western Cape of South Africa.

## Supporting information

**S1 Fig. Dynamic AHS prevalence in the KNP plain zebra population over the study period and stratified by age classes.** Temporal variation in the proportion of foals (Y), juveniles (J), and adults (A) that are infectious following 40 independent incursion events occurring on January 1st 2012. Here is shown the prevalence one year after the initial introduction event, assumed representative to the endemic circulation of AHS in the KNP population of plain zebra. Note the difference of scales in the *y*-axis.

**S2 Fig. Change in the risk of AHS persistence in the Western Cape Province (WCP) as a function of the timing of the introduction.** Distribution of the duration of the outbreaks generated from 40 independent incursion events occurring at the first day of each month between January 2012 and December 2015. Here we considered incursions occurred in the largest population of plains zebra reported in the WCP (N=168) at the start of each simulation. Horizontal black bars show the mean outbreak duration.

**S3 Fig. Spatial distribution of the risk of AHS persistence among all holdings with plains zebra in the Western Cape Province.** Risk of persistence was computed using either the median outbreak duration and the probabilities that AHS outbreaks would persist for >1 year, >2 years and >3 years. Risk measures were computed over 40 independent AHS introductions in all individual holdings. Here AHS was consistently introduced in April 2012.

**S4 Fig. Comparison of the daily variations in sunset temperatures between climatic zones A and B.** Daily variations in sunset temperatures for all sites considered in this work and driving holdings in (A) group A and (B) group B as defined in Fig 6.

**S5 Fig. Distribution of the risk of AHS persistence for increasing potential size of a plains zebra population in the WCP.** Risk of persistence was computed using either the median outbreak duration and the probabilities that AHS outbreaks would persist for >1 year, >2 years and >3 years. Risk measures were computed over 40 independent AHS introductions in considered scenarios. AHS was consistently introduced in April 2012 and considering an initial population density consistent to the median density reported in the WCP. Here is shown the outcomes for incursions in a typical holding located in either climatic zone A or climatic zone B.

**S6 Fig. Distribution of the risk of AHS persistence for increasing potential mean infectious period** 1*/γ* **of plains zebra and for various population sizes in the WCP.** Risk of persistence was computed using either the medium outbreak duration and the probabilities that AHS outbreaks would persist for >1 year, >2 years and >3 years. Risk measures were computed over 40 independent AHS introductions in considered scenarios. AHS was consistently introduced in April 2012 and considering an initial population density consistent to the median density reported in the WCP. Here is shown the outcomes for incursions in a typical holding located in either climatic zone A or climatic zone B.

## Acknowledgments

The authors gratefully acknowledge CapeNature for providing data and intellectual inputs on the zebra populations in the Western Cape Province. We also gratefully acknowledge Dr Stella Mazeri (Roslin Institute) for the compilation of the climate data and Dr Natascha Meunier (Roslin Institute) for many helpful discussions during the preparation of this manuscript. The authors also thank Dr Gert Venter (Agricultural Research Council – Onderstepoort Veterinary Research) for providing useful comments on the manuscript.

